# Microbial biogeography and ecology of the mouth and implications for periodontal diseases

**DOI:** 10.1101/541052

**Authors:** Diana M. Proctor, Katie M. Shelef, Antonio Gonzalez, Clara L. Davis Long, Les Dethlefsen, Adam Burns, Peter M. Loomer, Gary C. Armitage, Mark I. Ryder, Meredith E. Millman, Rob Knight, Susan P. Holmes, David A. Relman

## Abstract

Human-associated microbial communities differ in composition among body sites and between habitats within a site. Patterns of variation in the distribution of organisms across time and space is referred to as ‘biogeography’. The human oral cavity is a critical observatory for exploring microbial biogeography because it is spatially structured, easily accessible, and its microbiota has been linked to the promotion of both health and disease. The biogeographic features of microbial communities residing in spatially distinct but ecologically similar environments on the human body, including the subgingival crevice, have not yet been adequately explored. The purpose of this paper is twofold. First, we seek to provide the dental community with a primer on biogeographic theory, highlighting its relevance to the study of the human oral cavity. For this reason, we summarize what is known about the biogeographic variation of dental caries and periodontitis and postulate as to how this may be driven by spatial patterning in oral microbial community composition and structure. Second, we present a number of methods that investigators can use to test specific hypotheses using biogeographic theory.

To anchor our discussion, we apply each method to a case study and examine the spatial variation of the human subgingival microbiota of 2 individuals. Our case study suggests that subgingival communities in the aggregate may conform to an anterior-to-posterior gradient in community composition. The gradient appears to be structured both by deterministic and non-deterministic processes, though additional work is needed to test and confirm specific hypotheses. A better understanding of biogeographic patterns and processes will advance our understanding of ways to optimize the efficacy of dental interventions targeting the oral microbiota.

## Introduction

Different teeth and tooth aspects display differential susceptibility to caries, gingivitis and periodontitis^1–4^. Given that dental pathology arises in large part due to disturbances in microbial community membership, structure and/or function, these observations emphasize the utility of disentangling the relative effects of geographic (tooth location within the oral cavity and tooth surface location), environmental (such as tooth shape, size, morphology, etc.) and host-specific (genetic, socioeconomic, demographic, behavioral) factors on the structure and function of microbial communities in the oral cavity. Biogeographers have two aims: to describe the distribution of organisms across time and space and to identify underlying causal mechanisms that drive or maintain the observed patterns of heterogeneity^5^. Antony Philips van Leeuwenhoek conducted the first biogeographic survey of the organisms that inhabit the human oral cavity and published it in the year 1683, over 3 centuries ago^6^. Using microscopy, Leeuwenhoek observed key differences in the variety of bacteria found in saliva compared to tooth surfaces when comparing cells using discriminating characters such as cell shape, cell size, abundance, and motility. Since those pioneering studies, most biogeographic surveys of the bacteria that live in the human oral cavity have drawn the same conclusion: major morphological and molecular features distinguish organisms that have differing proclivities for different intra-oral habitats.

Though one of the earliest 16S rRNA gene-based surveys of microbial biogeography across human body sites showed that the features of microbial community assembly, like the morphological features of cells, also differ by body site^7^, the number of studies applying ecological theory^8^ to study the microbiome, and in particular the oral microbiome, has been limited. For this reason, insights into the types of spatial structures formed by microbial communities, the spatial scales (e.g., the spatial extent) over which microbial populations vary, and the causal mechanisms that maintain the spatial heterogeneity of the human oral microbiota have been limited^9^. We propose that understanding ecological processes underlying the biogeography of dental disease will inform our understanding of what is necessary to maintain or restore oral health. Before reviewing spatial patterns from the vantage point of the dental chair, we begin with a primer on biogeographic theory, which can be used by researchers to anchor their studies of temporal and spatial variation of the oral microbiome. Next, we review our current understanding of spatial patterning in oral diseases, and in particular, in dental caries and chronic periodontitis and the associated microbiota. We discuss the limitations of current approaches for understanding the similarities and differences of microbial community composition, structure and function between and across different intra-oral habitats. Finally, we present several statistical and ecological techniques that can be applied to study biogeography in the human oral cavity. To illustrate the utility of these approaches for exploration of oral biogeography, we use a case study approach and present an analysis of previously unpublished data. To facilitate the use of these methods by others, we provide the R code and data used to generate figures as described in the section on data availability.

## A primer on biogeography: ecological pattern and process

Biogeography is the study of the distribution of organisms over space and time^10^. The major questions biogeographers ask include: What enables a species to live where it does, and what prevents it from colonizing other areas? What role do environmental variation, biological interactions, and historical events (e.g., colonization history, past environmental conditions, etc.) play in shaping distributions? To answer these questions, biogeographers look for patterns in the distribution of diversity and propose mechanism-based hypotheses that can be tested to determine the contribution of ecological processes to community assembly. These mechanisms can be grouped into the four basic processes of community ecology: selection, ecological drift, diversification, and dispersal^11,12^. In this section, we define these processes and highlight how they may generate spatial patterning in the oral cavity.

### Selection

Selection encompasses a set of ecological features that operate on several features of fitness – survival, growth rates, and reproduction, which differ among organisms and the outcomes of which govern community assembly^13,14^. In the context of the human oral cavity, these ecological features can be grouped into four broad categories. First, the oral cavity is constantly exposed to microbes from the environment through open mouth breathing, dietary intake, and person-to-person contact. Not all microbes that enter the oral cavity are capable of residing in it. Thus, certain host-associated mechanisms – including active and innate immunity – selectively filter microbes, preventing their colonization. This colonization resistance is a selective process. Second, the organisms that are capable of colonizing either shedding or non-shedding intra-oral surfaces are subject to broad scale features that create physical gradients in the intra-oral environment. Physical gradients of temperature, fluid velocity or pH give rise to gradual but continuous differences in community composition and diversity across the gradient, impacting observed community structure^15,16,17^. Third, other features give rise to patchy structures in microbial community diversity and composition in which each patch is separated by a discontinuity. For example, independent of the clinical features of a carious lesion it is possible to use the microbiota to distinguish between a carious and sound site^18–20^. Our ability to use the microbiota to distinguish between a carious lesion and a sound surface in this manner suggests dental caries is a patchy process. Finally, fine-scale biotic processes such as local species extinctions, reproduction, predator-prey dynamics, symbiotic interactions, etc. induce spatial patchiness in community structure – even in relatively homogenous communities, indicating that community structure is an emergent function of ecosystem processes^21^. Microbiome focused studies should endeavor to include measurements of factors thought to exert a selective effect on microbial communities, including intra-plaque pH and redox potential, immunological features, and salivary or gingival crevicular fluid flow rates, for example. These measurements are necessary for understanding the high dimensional data generated by modern ‘omics’ technologies, including RNASeq, metagenomics, 16S rRNA gene profiling, and metabolomics.

### Dispersal

Dispersal describes the movement of organisms across time and space. Microbial dispersal is accomplished by both passive motility (e.g. transport in fluid flow) and active motility, including chemotaxis. Whether the mechanism is passive or active, dispersal incurs costs to the organism. Active dispersal costs energy through the consumption of cellular resources^22^ while all dispersing organisms encounter some level of risk, particularly those that are not using chemotaxis: dispersal means that an organism leaves a patch where reproduction empirically occurred in search of another location where conditions may be less favorable^23^. At least two distinct features characterize dispersal in the human oral cavity. First, the human oral cavity is seeded from the mother and other caregivers at birth and during infancy^24^. The oral cavity is open and continuously exposed to environmental microbes from other individuals, the air and from the rest of the environment. Second, dispersal may occur among sites within the oral cavity. Organisms may additionally disperse across teeth or between sites (e.g., from the teeth into saliva).

Dispersal may explain results of studies focused on scaling and root planing (SRP), an intervention that treats sites impacted by periodontitis. For example, multi-locus sequence analysis revealed clearance of *Porphyromonas gingivalis* for most patients after treatment. However, 3 of 12 patients who were colonized prior to treatment were again colonized by the same strain at some point after treatment^25^. This suggests that reservoir sites external to the site of intervention may have served as the source of reinfection. Indeed, many studies evaluating the efficacy of a single SRP treatment to eliminate periodontal pathogens from an affected pocket have demonstrated that communities revert to the pre-cleaning state after a temporary reduction in pathogen load^26,27^. Since the size and density of the source population is a determinant of dispersal, controlling the oral load of each pathogen (i.e., controlling the population size of potential colonists) may prove important in managing periodontitis. Indeed, repeated bouts of SRP with maintenance therapy more effectively reduces the load of bacterial pathogens than single bouts of SRP^28^. Studies that focus on modeling the source population size and the geographic distance between sites may be able to identify areas of weakness where a treatment approach is failing. Modeling approaches, in general, are likely to provide new insight into the role that dispersal plays in structuring communities.

### Drift and Diversification

Drift occurs as a consequence of non-deterministic differences in demographic features (e.g., birth, death and reproduction) between populations. This random demographic variability will differ among sites irrespective of the surrounding environment, leading to spatial heterogeneity in community diversity and composition^10^. Importantly, the interaction of drift and any of the other ecological processes may have important impacts on community dynamics. As discussed above, a large source microbial population is associated with high rates of dispersal. In the context of disturbance, drift and dispersal can interact and elevate the role of drift on community assembly. For example, a taxon that is present at low abundance in a community prior to a disturbance can come to dominate that community if it survives – due to chance events alone – the disturbance at an abundance that permits it to outcompete other colonists during community re-assembly^11^. The interaction between drift and dispersal thus has important implications for clinical interventions such as SRP that non-selectively reduce community biomass.

Ecological diversification represents a balance between speciation (i.e., when a population splits into a pair of lineages) and extinction (i.e., the loss of a lineage due to its elimination in the niche)^29^. In the absence of either selection or dispersal, a single species occupying two spatially segregated sites may experience the same environment and yet diverge into more than one lineage due to the accumulation of different random mutations in different individuals at each site^14^. In addition to random mutations, recombination and horizontal gene transfer can give rise to diversification in species^30,31^, including the inhabitants of the oral cavity. Since the same random event rarely occurs twice, such a process can give rise to heterogeneity in community structure that is independent of environmental selection^10,12^.

## Spatial patterns of dental disease

Complex systems are likely structured by more than one of these four ecological processes of selection, dispersal, drift and diversification. Studies that seek to assess patterns of spatial variation in the oral cavity should, during the experimental design phase of the study, consider strategies that enable the testing of specific processes. Our ability to understand the relative contribution of these four ecological processes to intra-oral microbial community structure will help explain why current clinical interventions work, when and why they fail, and lead the way to new and improved therapies. Here, we set the stage for further discussion of these processes by reviewing what is known about the spatial patterning of dental disease and the tooth-associated microbiota.

### Spatial patterns of dental caries and the supragingival microbiota

Epidemiological studies consistently reveal a spatial pattern for dental diseases including dental caries. For example, the occlusal^3,32^ and proximal^33^ surfaces of the first molars in both jaws are more frequently affected by dental caries than any other surface in the mouth. Most explain these patterns by observing that the pits and fissures of those teeth are relatively hidden from the protective activity of the tongue, toothbrushes and salivary flow. But why would the buccal surface of the lower first molar be the most susceptible of all *buccal* surfaces to cervical caries^3,34,35^? What processes explain the generally low incidence of dental caries on the canines and incisors of healthy individuals^3,32–34,36–38^? And then why would the incisors and canines be the second most frequent site of caries in an experimental model in man^1^? Striking shifts in the spatial patterning of dental disease imply the occurrence of a corresponding and antecedent shift in the spatial pattern of the tooth-associated microbiota since it is aberrant composition, structure and function of the microbiota at the tooth surface that gives rise to the pathology associated with dental caries^39^.

Environmental selection of spatially segregated microbiota offers an explanation for the spatial patterning of dental caries. The velocity of the salivary film flowing over individual tooth surfaces varies based on tooth position. Elegant work from the Colin Dawes group demonstrated that the velocity of the film flowing over the buccal surface of the lower first molar is 1 mm/min, and the rate of salivary clearance is 44.8 minutes^40–42^. On the other hand, the velocity of the salivary film flowing over the buccal surface of the upper 1st molar is 4.6 mm/min, and the rate of salivary clearance is 3.5 times faster (12.6 min). These differences in salivary film velocity and salivary clearance have direct implications for the physiology and the ecology of the dental plaque communities. The buccal surface of the lower first molars experience a more profound drop in plaque pH than those of the upper first molars following a glucose challenge^43^, suggesting that the indigenous inhabitants of the buccal surfaces of the lower molar have greater metabolic potential than the corresponding communities in the upper jaw. Thus, environmental selection may provide an explanation for the observation that the buccal surface of the lower first molar is the most frequent site of root caries in experimentally “de-salivated” rats fed a cariogenic diet^44^, and could partially explain why the mandibular molars have the highest root caries increment (RCI) across all ages^45^.

When the salivary glands are not stimulated, the lower lingual region of the mouth has the highest salivary film velocity (7.8 mm/min), as well as the shortest clearance half time (8.7 min), as might be expected from this region’s proximity to the submandibular and sublingual glands.

In contrast, the upper anterior buccal region of the maxilla, which may be bathed primarily by labial gland secretions, has the lowest film velocity (0.8 mm/min), and the longest clearance half time (70.2 min) of all oral compartments. When sucrose containing gum (but not a lemon drop) is used to stimulate salivary flow, the estimated flow rate approaches that of a well-mixed solution, and the clearance half time drops considerably for the lower lingual incisors, lower lingual molars, and the buccal upper molar regions^46^, but not for other sites. Restricting salivary flow experimentally causes intra-plaque pH not only to drop to levels that tip the balance towards dental enamel demineralization, but it also reduces salivary clearance of sugars and acids^15^. Geographic variability in the volume of saliva distributed across soft tissue sites has also been demonstrated^47,48^. Taken together, these studies suggest that under normal circumstances there may be differences in plaque pH at different oral locations in the mouth, even in healthy individuals. Such site-to-site heterogeneities presumably not only imply that certain dental surfaces are more or less susceptible to demineralization, but they also provide a primary basis (e.g., pH) for structuring the biogeography of the oral microbiota. This raises the question as to whether reduced salivary flow alters the spatial structure of oral microbial communities.

Most prior studies that have evaluated the microbiota of individuals experiencing significant, chronic reductions in salivary flow (known as hyposalivation) have done so largely without regard for habitat structure in the mouth; most have used rinsing samples^49–55^. Other work only sampled a single supragingival surface^56^. Of the studies that did sample multiple sites, such as the maxillary and mandibular molars, samples were pooled before analysis^57–59^, or summary data were reported for whole mouths rather than site-specific data^60,61^. To our knowledge the first study to examine spatial variability (two interproximal sites, anterior vs. posterior) in cultivable *Lactobacillus* populations, did so by studying Sjögren syndrome patients, and provided the first evidence in support of the hypothesis that reduced salivary flow remodels the spatial organization of oral microbial communities^62^. The paucity of studies examining site-to-site variability of the oral microbiome is not restricted to research on the impact of hyposalivation on oral microbial communities. Investigation of the spatial organization of intra-oral communities with normal salivary flow is also lacking.

Recent imaging studies have identified several complex structures in microbial biofilms in the human oral cavity, highlighting spatial variation at the micron-scale^63^. Only a handful of studies have examined the broad scale spatial pattern of communities between and across teeth^16,64–66^. The first seminal work exploring spatial variation of supragingival communities found “tooth number” to be significantly associated with variation in the proportions of 20 out of 40 taxa, even after accounting for differences in the biomass of the communities found at different sites^64^. Other work reported that microbial communities clustered by tooth class as well as by tooth surface type, suggesting consistent patterns of spatial heterogeneity^65,66^.

We recently incorporated geographic coordinates into an explicit spatial model analysis of supragingival communities^16^. In that work, we reported that supragingival and soft tissue microbial communities vary along an anterior-to-posterior gradient in the mouths of healthy individuals. Importantly, the ecological gradient appeared to be attenuated in individuals with low salivary flow due to Sjögren’s syndrome. Future work is required to disentangle the relative effect of salivary film velocity from that of dispersal limitation due to geographic separation, tooth morphology, and other factors. Nonetheless, extant studies definitively provide evidence for spatial patterning in the distribution of dental caries and provide early support for the hypothesis that supragingival communities are geographically structured, which may explain spatial patterning in dental caries.

### Spatial patterns of periodontal disease and the subgingival microbiota

Spatial patterning has also been observed in gingivitis and periodontitis. One of the earliest studies of experimental gingivitis in man revealed the uniform development of subgingival plaque throughout the dentition in the oral cavities of 12 dental students who deliberately discontinued oral hygiene^2^. Intriguingly, no differences were found in plaque accumulation between different tooth classes or when comparing all tooth aspects (buccal, interproximal, lingual). Lingual tooth surfaces, on the other hand, could be distinguished by a paucity of debris accumulation compared to buccal and proximal sites. This observation was thought to be explained by the effect of the movements of the tongue, which are primarily geographically restricted to the lingual tooth surfaces. Where plaque did occur, Löe et al. hypothesized that the age of the plaque contributed to its development such that the developing community altered the local environment in a manner that was permissive for the growth of organisms not typically found in health. Thus, there may be an interaction between environmental and biotic processes in the transition between states of oral health and disease. This hypothesis is now known as the ecological plaque hypothesis and is the prevailing explanation for the occurrence of dental _disease_^39,67^.

Here, we suggest a critical need to incorporate explicit spatial analysis of the microbiota into future tests of spatial patterning in dental disease. To understand disease, these patterns and processes in the oral microbiota must be linked on a site-specific basis. Prior work has suggested that periodontal disease displays left-right contralateral symmetry within individual mouths^4,68^. More recently, the Arteriolosclerosis Risk in Communities study revealed that a single tooth with periodontal pockets of at least 4 mm is highly predictive of that tooth’s contralateral pair experiencing a pocket of at least that depth^69^. Furthermore, various features of periodontitis (e.g., attachment loss, bleeding on probing, calculus deposition) appear to differ by jaw and tooth class. In China, the 4^th^ National Oral Health Survey of over 100,000 adolescents and another study of 398 adults revealed significant patterns of symmetry in gingival bleeding in addition to a difference between jaws in the distribution of pocket depths and attachment loss^70,71^. An investigation of periodontal health in over 11,000 adolescents in the United States determined that gingival bleeding, attachment loss and calculus deposition occur to similar degrees at symmetric sites in the mouth, and most often at molar sites in the maxilla and incisors in the mandible and less frequently impacting the bicuspids of either jaw^72^. Other reports similarly confirm significant effects of tooth class and jaw in the spatial patterning of periodontitis^71,73,74^.

What might explain these patterns of periodontal disease? In addition to elements of interpersonal variability, including host genetics, comorbid medical conditions, social determinants, and lifestyle and behavioral habits, the local tooth environment likely contributes to the spatial patterning of dental disease^75,76^, which is mediated in part by the microbiota. Changes to the subgingival habitat in gingivitis, the mildest and most common form of periodontal disease, include increased inflammation and gingival crevicular fluid (GCF) flow without clinical loss of the of the supporting surrounding tissue^77–79^. As the connective tissue is destroyed by microbe-induced inflammation, epithelium from the dentogingival interface migrates in an apical direction along the root surface. The formation of periodontal pockets provides an increased surface area for microbial colonization along the tooth surface, resulting in a marked increase in microbial load between periodontal health and periodontitis^80–82^. Estimates of bacterial biomass in health are around 10^3^ – 10^6^ bacterial cells per mL with an average probing depth of 1.8-mm^67^ while sites with probing depths of 4-12 mm may harbor anywhere from 10^7^ to 10^9^ bacterial cells per mL^78,83^. Differences in biomass may prove important in the ecology of the subgingival crevice since the likelihood that dispersal will occur depends in part on the size of a population.

Importantly, symmetric patterns of periodontal disease appear to correspond to symmetric variation in the composition of the microbiota. Sites that are culture-positive for *Prevotella intermedia, Prevotella nigrescens* or *Aggregatibacter (Actinobacillus) actinomycetemcomitans* at a site on one side of the mouth are highly predictive of an increased burden for those same organisms at a symmetrical site^4^. Other work reported sporadic colonization of *A. actinomycetemcomitans* at spatially segregated sites in 2 healthy individuals with generalized periodontitis^84^. In that work, *Porphyromonas gingivalis* colonized a greater number of sites compared to *A. actinomycetemcomitans,* indicating possible differences in species ranges or sample size limitations. A second spatial pattern was observed in the distribution of *Prevotella gingivalis* and *P. intermedia,* both of which were more abundant at posterior compared to anterior sites^85^. Further support for spatial segregation of subgingival communities across the anterior-to-posterior dimension comes from studies reporting an anterior-to-posterior gradient in subgingival temperature, which was associated with differences in microbial colonists in the subgingival crevice^17^. Finally, a variety of taxa were found to preferentially colonize different tooth classes^86^ while other work suggests communities in the subgingival crevice differ on opposing aspects of teeth, even when comparing different facets of a single tooth^66^. Thus, spatial patterns of microbial community composition may explain spatial patterns in the occurrence of gingivitis and periodontitis. Future work should integrate the investigation of epidemiological, environmental and microbiological correlates to solidify our understanding of the relationship between patterns in the microbiota and patterns in dental disease.

## Methods for investigating biogeographic patterns

A challenge for oral microbial ecologists is the identification of the types of spatial patterns present in the oral microbiota, the processes underlying the patterns, and how they pertain to disease. Many extant studies of the oral ecosystem use saliva, rinsing samples, or pooled plaque samples to examine the composition and diversity of intra-oral microbes^87–92^. These broad scale sample collection methods do not permit analysis of spatial pattern and process and therefore limit our understanding of dental disease. Investigation of the processes underlying the spatial patterning of dental caries or periodontitis requires high resolution sample collection schemes that include multiple teeth and tooth aspects per human subject. Such sample collection schemes can easily generate terabytes of sequencing data for thousands to tens of thousands of samples^16^, making data analysis a challenge. In this section, we provide an overview of statistical and ecological methods that can be used to explore spatial patterns and processes. To illustrate the utility of these approaches we employ a case study approach, analyzing subgingival samples collected from the mesiobuccal aspect of all teeth (excluding 3^rd^ molars) of 2 medically and dentally healthy individuals. A complete description of the patients sampled, the data collected and the methods used to analyze the data have been previously described^93^. The purpose of this case study is to describe techniques in a manner that highlights their utility in the applied analysis of the oral microbiome.

### Exploratory analysis of spatial patterns

Exploratory data analysis is often the first step in a data analysis pipeline. Here, we illustrate two approaches to visualizing spatial patterns in geographically structured systems. First, many researchers seek to define the role that a specific pathogen plays in initiating or sustaining oral disease. Researchers interested in understanding the spatial distribution of one or more pathogens might start by visualizing the distribution of each pathogen across the sample sites that were surveyed, such as at the lingual or buccal tooth aspect. We have previously published R code that may be used by others to visualize such spatial patterns in various contexts^16^.

To demonstrate the power of visualizing taxonomic distributions across sites in the context of the oral cavity, we used SitePainter to examine the distribution of 2 randomly selected genera – *Prevotella* and *Fusobacterium* – at the subgingival surfaces across the mesial-buccal aspect of all teeth (excluding 3^rd^ molars) and the mesial-lingual aspect of 2 teeth (14, 15) within each of 2 individuals (**Figure 1**).

**Figure 1:**
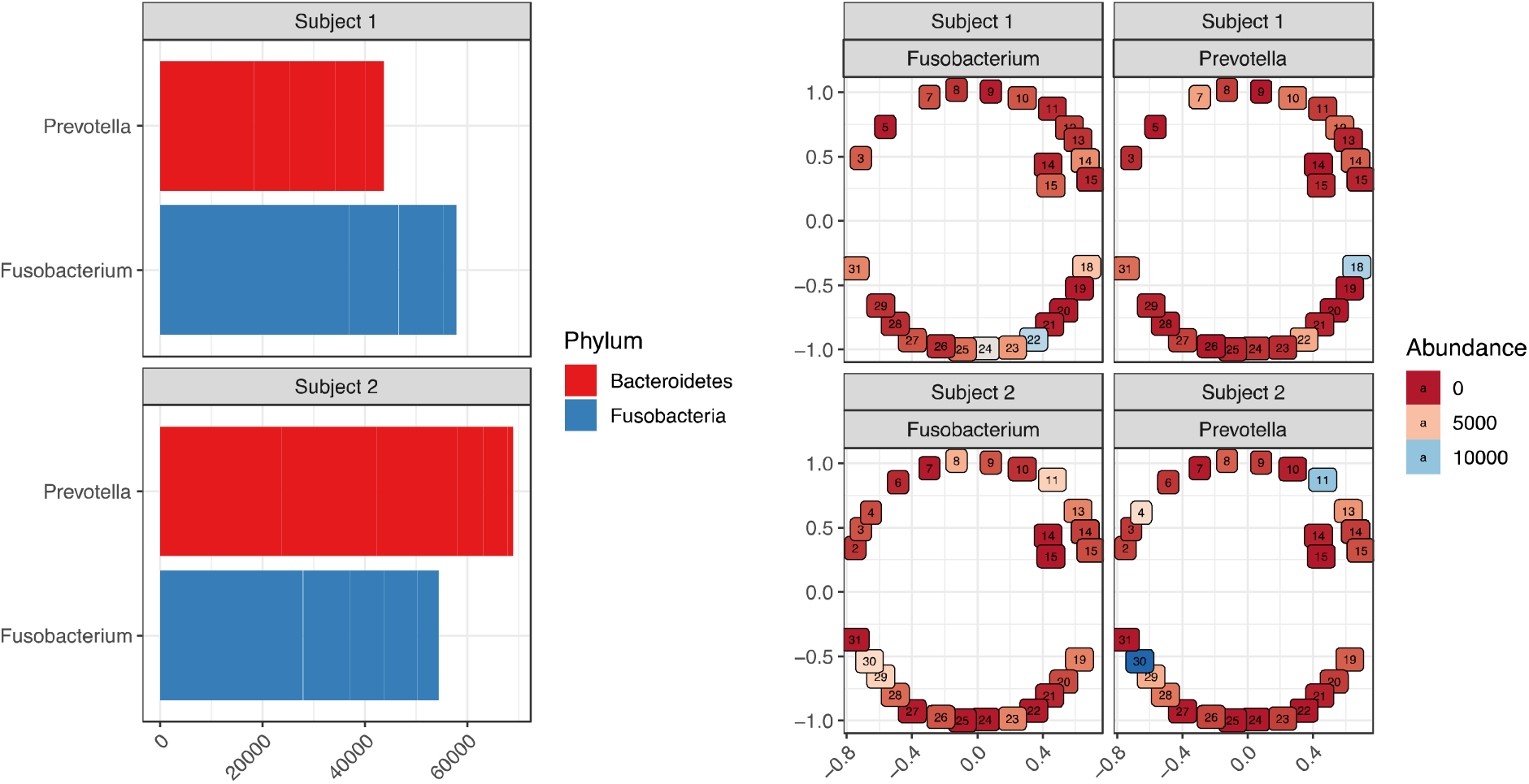
Exploratory analysis of 4 genera. a) The absolute abundance (x-axis) of *Prevotella* and *Fusobacterium* is plotted (y-axis) for each of 2 individuals. Each bar is shaded by the Phylum to which the taxon belongs. b) the site-specific abundance of each Genus was visualized for each site for Subject 1 (Left) and Subject 2 (Right). Similar colors represent similar abundance values: red indicates low; pink intermediate; and blue high abundance values for each taxon at that site. Numbers within each circle correspond to universal tooth number.

In this example, *Fusobacterium* appears to be present at roughly comparable abundance levels in both subject 1 and 2 (**Figure 1A**). Therefore, approaches that pull out interesting taxa based on total differential abundance^95^ between individuals would fail to flag *Fusobacterium* as an interesting feature. One might ask, however, whether *Fusobacterium* colonizes all teeth in roughly equal proportions in one or both subjects or whether it exhibits some degree of site-specificity that is concordant between subjects. Examining the spatial distribution of *Fusobacterium* across mouths reveals subject-specific patterns of colonization and an absence of site-specificity (**Figure 1B**). Specifically, Subject 1 has the highest *Fusobacterium* colonization at teeth 18, 22, 23, and 24 while Subject 2 has the highest *Fusobacterium* colonization at teeth 8, 11, 29, and 30. *Prevotella* similarly exhibited subject-specific differences in patterns of colonization and a lack of concordance between impacted sites. If generalized across a larger number of subjects, these observations would suggest that colonization of individual tooth surfaces by these two genera occurs randomly and that these taxa tend to lack a predilection for the anterior vs. the posterior compartment or even for any specific tooth class.

For researchers who are not comfortable using R to visualize data, a utility that obviates the need for familiarity with R and which was developed for visualizing spatial patterns is the tool SitePainter^94^.

A more rigorous and less exploratory approach would be to query the data to identify interesting taxa that vary in their abundance across teeth. Researchers who hope to identify spatially variant taxa may calculate coefficients of autocorrelation such as Moran’s I for each taxon in the dataset^96^. This statistic when calculated for each taxon identifies those that vary as a function of the geometric distance, which can be defined for example as the Euclidean or Manhattan distance, separating sample sites. The coefficient can be either positive or negative, small or large, and is associated with a p-value. Researchers can select the subset of taxa that appear to be spatially variant by picking taxa that meet certain threshold p-values and coefficient sizes. The spatial distribution of these selected taxa can then be visualized using SitePainter or various software packages in R to examine the specific pattern underlying the summary statistics. The application of Moran’s I in this context is univariate in nature (i.e., one coefficient per taxon) and p-values should be corrected for multiple testing.

### Statistical approaches to identifying spatial patterns

A variety of well recognized statistical methods and models have been developed to identify spatial patterns in complex communities^21,97^. In this section, we describe 3 statistical approaches – trend surface analysis, principal components analysis of neighbor matrices (PCNM) and Moran’s Eigenvector Maps – that can be used to uncover multivariate spatial patterns in complex communities^96,98^. By conducting a study that encompasses all 3 methods it becomes possible to evaluate the robustness of any given pattern as well as its spatial extent.

Trend surface analysis is a multivariate method that explicitly considers the geographic coordinates of sample sites. The geographic coordinates are used to construct an orthogonal 2^nd^ or 3^rd^ degree polynomial function of the geographic coordinates of sample sites. This polynomial function is then used as a predictor in a principal components analysis with respect to instrumental variables (PCA-IV). In effect, the ordination is constrained by the geographic coordinates of sample sites. This method generally identifies broad scale spatial patterns such as large-scale ecological gradients.

Principal Components Analysis of Neighbor Matrices (PCNM) is a method similar to trend surface analysis in that it indirectly uses geographic coordinates as a constraint in an ordination. Unlike trend surface analysis, several features permit PCNM to detect both broad and fine scale spatial patterns^21^. Rather than using a polynomial function as a constraint in a PCA-IV, PCNM uses a neighborhood distance matrix which is constructed in three steps. First, the geographic coordinates of sample sites are used to construct a geographic distance matrix (e.g., Euclidean distance, Manhattan distance, etc.) that describes all pairwise distances between sample sites. Second, a neighborhood matrix is constructed by defining as neighbors any site within an arbitrarily defined geographic distance threshold. The distance threshold that is used to define the network thus determines the spatial scale that is examined (i.e., whether it is a broad or fine scale). Finally, the neighborhood matrix is subject to a principal coordinates analysis; the resulting eigenvectors are used as “PCNM variables” (i.e., as spatial predictors) in a redundancy analysis (RDA). The RDA will thus maximize the variance in community composition across a linear combination of the PCNM variables. The investigator may analyze the results as with any other ordination. In addition to identifying the ordination axes that are significant it is possible to use permutation testing to identify the PCNM variables that seem to correlate with the significant axes.

One feature of PCNM that can be problematic is its reliance on an arbitrary cutoff to define the neighborhood. At the current time, calibration experiments are needed to define an appropriate threshold to model microbial dynamics in the human oral cavity. Moran’s Eigenvector Maps (MEM) provides another avenue to explore spatial patterning in a manner similar to PCNM but without the need for an arbitrary threshold. Rather than defining a single neighborhood matrix, MEM defines multiple different spatial models each differing from the other by its neighborhood truncation threshold. PCA-IV or RDA is performed on each model and the optimal model is selected using a model selection criterion such as Akaike information criterion (AIC). Using this method, it becomes possible to define the distance threshold that best describes a microbial neighborhood in the oral cavity. In summary, trend surface analysis identifies broad-scale spatial patterns while PCNM and MEM can identify both broad and fine-scale patterns.

#### Case Study – applying statistical models to the subgingival microbiota

We previously used trend surface analysis, PCNM, and MEM to demonstrate that microbial communities inhabiting a diverse set of tissues in the oral cavity conform to an anterior-to-posterior gradient^16^. Here, we illustrate the application of trend surface analysis and MEM to the analysis of communities in the subgingival crevice.

First, we performed a trend surface analysis on the dataset for the 2 subjects for whom subgingival samples were collected. To visualize the spatial pattern, we plotted the first principal component as a function of tooth number, plotting each subject individually in distinct panels (**Figure 2A**). Generally speaking, molars (blue) tended to share negative scores along axis 1 while the incisors (green) shared positive axis 1 scores. The trend line indicates a gradual and continuous change in community composition across the anterior-posterior dimension. Comparing the two subjects to each other revealed marked inter-individual variation in the degree to which samples separate across the anterior-to-posterior dimension. For both individuals surveyed, the relative difference between the molars and the incisors appeared to be greater in the mandible (tooth 16-32) compared to the maxilla (tooth 1-15) while the absolute difference between the molars and incisors appeared to be greater in subject 1 compared to subject 2. Thus, the trend surface analysis identifies an ecological gradient in community composition between the molars and the incisors which appears to be variable when comparing individuals.

**Figure 2:**
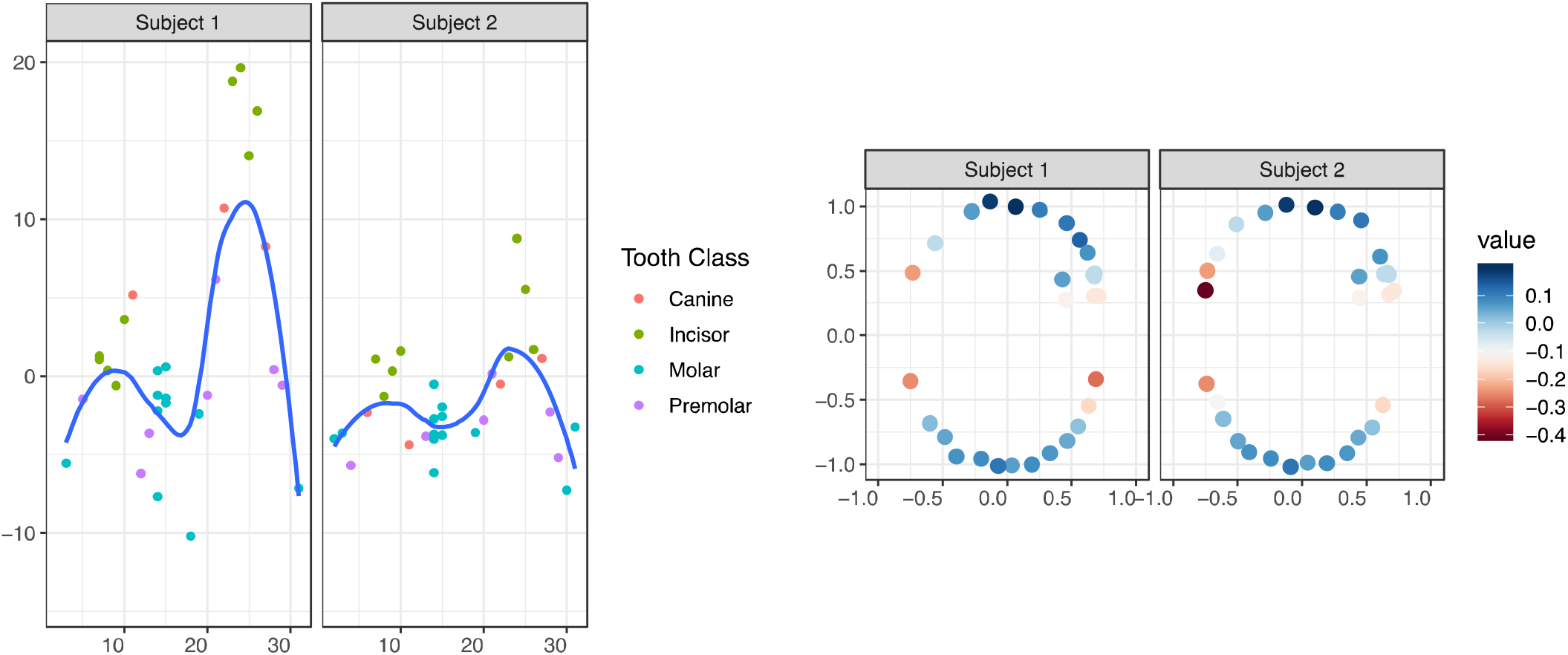
Subgingival communities, like supragingival communities, may conform to an ecological gradient. a) Trend surface analysis was used to examine spatial patterning at subgingival sites. Scores from the first principal component (y-axis) is plotted against universal tooth number (x-axis). Each point represents a sample that is colored according to tooth class (canine, incisor, molar, premolar). The blue line is a loess smoothened curve surrounded by 95% confidence intervals in grey. b) MEM was used to evaluate spatial patterning in subgingival samples from the same subjects. Each point represents a tooth plotted against the x-and y-geographic coordinates of sample sites. Points are shaded with a heatmap scale according the first RDA axis. The trend surface and MEM models both suggest communities vary along an ecological gradient that distinguishes between sites across the anterior-to-posterior dimension.

Next, we sought to evaluate the broad and fine-scale spatial patterns in community composition using Moran’s Eigenvector Maps (MEM). We generated 20 different geographic distance matrices each truncated randomly at a different distance threshold. The optimal MEM model was identified as the one that minimized Akaike information criterion (AIC). The optimal model corresponded to a Euclidean distance threshold of 0.89 and explained approximately 4% of the total variation in the data. A permutation test revealed that the first and only the first ordination axis explained a significant fraction of the variation in the data (F=2.4, p=0.002). Further, only one spatial predictor explained a significant fraction of the variation in the data (F=2.4, p=0.004). To examine the significant spatial structure, the first ordination axis was projected onto the x and y coordinates of the subgingival sample sites (**Figure 2B**). Teeth in the anterior mouth tended towards positive scores along the significant ordination axis while the posterior teeth tended towards neutral to negative scores along that axis.

Trend surface analysis uses a smoothened polynomial function as a predictor while MEM uses eigenvectors derived from decomposing a constellation of truncated geographic distance matrices as a predictor. Despite the slight differences in each approach, they both identified similar features in community variation across sites in the subgingival crevice, suggesting that this broad scale ecological gradient in the subgingival crevice may be robust to further inquiry.

### Ecological approaches to identify spatial patterns

Statistical approaches can be used with ecological models in order to evaluate the robustness of a given spatial pattern. The statistical approaches described so far explicitly employ the geographic coordinates of sample sites to model spatial patterns. One advantage of some ecological models, on the other hand, is that they can identify spatial patterns without explicit geographic modeling. A second advantage is that specific spatial patterns can be identified through the elements of metacommunity structure (EMS) approach. EMS models distinguish between 6 patterns in geographically structured communities, including 1) nested subsets, 2) checkerboards, 3) Clementsian gradients, 4) Gleasonian gradients, 5) evenly spaced gradients and 6) random distributions^99–101^. Before defining these types of spatial patterns, we must first define several features of communities – coherence, turnover, and clumping, which distinguish between each of the spatial patterns.

Coherence assesses the extent to which species ranges across a gradient overlap across sites. Coherence is usually evaluated by examining an incidence matrix in which site occupancy by a given species is denoted by a ‘1’ and absence by ‘0’. Completely coherent species ranges occur when a species occupies all sites without any absences across the range. For example, in a model community where one species occupies 5 sites, a completely coherent species range would be defined in the incidence matrix as [1, 1, 1, 1, 1] since the species occupies all sites. Based on this definition, patterns such as [0, 1, 1, 1, 1] or [0, 1, 1, 1, 0] or [0, 1, 1, 0, 0, 0] would also be coherent. ‘Embedded absences’ are defined as interruptions in site occupancy and would be indicated by the pattern [0, 1, 1, 0, 1] in an incidence matrix.

In assessing a community, the EMS model organizes the incidence matrix in such a way that maximizes the coherence of all species in the community across sites. Coherence decreases as the number of embedded absences increases. Coherence is usually assessed by comparing the observed number of embedded absences to that observed in a null model, generated by simulating a set of random matrices. ‘Negative Coherence” occurs when the number of observed embedded absences is significantly greater than the number of embedded absences predicted by a random model. Negative coherence suggests that a community fits a checkerboard pattern in which pairs of species competitively exclude each other at sites across their ranges. ‘Positive coherence’ on the other hand suggests communities are structured along at least one common gradient. When the number of embedded absences is significantly less than the predicted number under a random model, communities are said to exhibit ‘positive coherence’.

In cases where positive coherence is observed it is possible to examine the specific type of gradient in community composition by examining two other metrics, turnover and boundary clumping. Turnover is the number of times that a species is observed to replace another when moving from one site to another. Observed turnover can be compared to the expected turnover calculated as the mean of a set of random community matrices. A nested subset describes a community that exhibits coherence but experiences less turnover than is expected, by chance, across sites^102^. The type of gradient can be further specified by examining the ‘boundary clumping’ metric, which measures the degree to which the boundaries of different species ranges cluster. Species loss can occur in a clumped, random, or a hyper-dispersed manner. The Boundary Clumping metric is assessed with the Morisita index. When the Morisita index exceeds 1, the boundaries of species ranges tend to be more clumped than expected by chance. On the other hand, a Morisita index that is significantly less than 1 indicates that less boundary clumping is observed across sites than expected.

As formally reviewed elsewhere^99^, spatial patterns in community ecology can be defined in terms of coherence, turnover, and boundary clumping. A ‘checkerboard pattern’ describes communities that exhibit negative coherence. Checkerboard patterns occur due to the competitive exclusion of taxa from sites across space. A ‘nested subset’ refers to a pattern of positive coherence and negative turnover – different sites appear to be colonized by subsets of a set list of species. A ‘Clementsian gradient’ exhibits positive coherence, positive turnover, and positive boundary clumping in which discrete communities replace each other across sites. A ‘Gleasonian gradient’ describes communities that exhibit positive coherence and positive turnover but random boundary clumping, i.e. species ranges along the gradient are random.

‘Evenly spaced gradients’ occur when there is positive coherence, positive turnover, and negative boundary clumping – discrete communities cannot be defined, but species appear to be ordered in a pattern that deviates from a random distribution. Finally, ‘Random distributions’ are defined by the absence of coherence, turnover, and boundary clumping.

#### Case Study – using EMS to identify spatial patterns

We used the EMS framework to identify the type of spatial pattern present in the subgingival crevice. Specifically, we computed coherence, turnover and boundary clumping for each of the two individuals in our case study (**Figure 3**). The observed number of embedded absences (embAbs) was higher for Subject 1 than for Subject 2 (**Figure 3A**). Regardless of subject, however, the predicted number of embedded absences under a random model (simMean) was significantly higher than the observed number of embedded absences across subgingival communities (p < 0.001). This observation is consistent with a spatial pattern of positive coherence and implies that subgingival communities are structured along at least one common ecological gradient. Further, this pattern supports the pattern detected using trend surface analysis and MEM (**Figure 2**).

**Figure 3:**
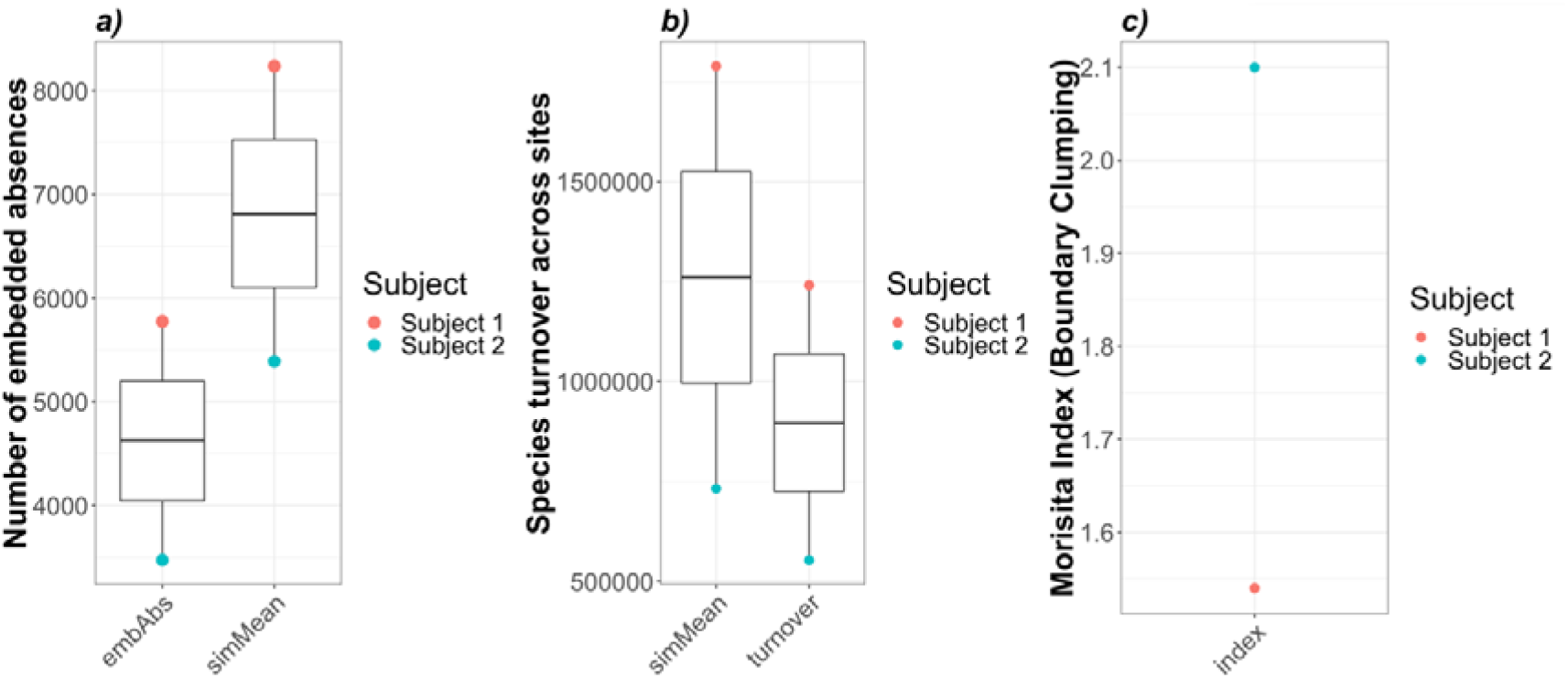
Elements of community structure suggests subgingival communities conform to a nested subset gradient with clumped species loss. a) coherence was less than the simulated mean for both subjects (p < 0.001). b) turnover was less than the simulated mean for both subjects (p < 0. 001), and c) boundary clumping exceeded 1 for both subjects. Taken together, these data suggest that the gradient fits a nested subset with clumped species loss.

Since the subgingival communities exhibited coherence we next examined turnover across sites (**Figure 3B**). Species turnover across sites was higher for subject 1 than for subject 2 but in both cases the observed turnover was higher than that predicted under a random model (simMean, p < 0.001), indicating that the specific spatial pattern may be described as a “nested subset”. This pattern implies that communities experience less turnover than would be expected along an environmental gradient. Next, we examined the degree of boundary clumping in the subgingival crevice (**Figure 3C**). Boundary clumping was higher for Subject 2 than for Subject 1. In both subjects, however, the Morisita Index exceeded 1, indicating that the communities are distinct and discrete (i.e., Clementsian) and that groups of species are lost at only one end of the gradient. Taken together, the EMS model implies that subgingival communities conform to a spatial gradient known as a nested subset with clumped species loss across the gradient. Intriguingly, we have previously reported this pattern for supragingival communities^16^.

By using both statistical and ecological models to examine the communities of the subgingival crevice it becomes possible to identify spatial patterns. Spatial patterns are emergent functional features of the ecosystem^11^. If coherence, turnover, and boundary clumping are shown to be important and common signs of health then the loss of these features might be subtle but important signs of early disease (the onset of which might not be otherwise known). As such, clinicians may be able to use spatial patterns to understand the extent to which the subgingival system is healthy or perturbed and the extent to which a treatment restores a perturbed ecosystem to its natural state.

### Understanding ecological processes underlying spatial patterns

The statistical and ecological methods described so far are able to identify spatial patterns, but they cannot elucidate the processes – selection, diversification, drift, dispersal – that drive those patterns. In the remaining sections we describe some of the approaches that can be used to test hypotheses about 3 ecological processes, drift, dispersal and selection.

#### Drift and the neutral model

The influence of stochastic, non-deterministic factors on community structure can be estimated by the application of a neutral model. Neutral theory holds that within-trophic level taxa are ecologically equivalent and that differences in distributions arise from the stochastic, demographic processes of birth, death, and migration (Sloan et al. 2006). The neutral model essentially acts as a null model that allows researchers to determine how much of the observed biogeographic pattern is due to stochastic differences among separate sites. The degree to which observed data fit neutral predictions varies among microbial assemblages. Relatively good fits have been observed for communities from freshwater lakes, healthy human lungs, tree-holes, wastewater treatment facilities, and deserts^103–107^. On the other hand, poor to conflicting fits have been observed for human gastrointestinal tract communities and coastal *Vibrio* populations^105,108–110^.

One benefit of the Neutral Community Model (NCM) is that it predicts how each individual taxon should be distributed across communities within the metacommunity. Deviations from this prediction identify taxa with interesting ecological characteristics. Taxa found more frequently than expected given their mean relative abundance in the metacommunity are likely to be under favorable selection by the environment and/or to have higher migration rates than their similarly abundant peers. Likewise, taxa found less frequently than expected given their abundance in the metacommunity are likely under negative selection at some sites and/or have lower migration rates than their similarly abundant peers.

##### Case Study – applying the neutral model to the subgingival microbiota

In order to identify ecological processes that may underlie spatial patterns in the subgingival microbiota we applied the NCM to the subgingival samples from our 2 subjects.

The frequency-abundance distribution of taxa conformed relatively well to the distribution predicted by the NCM. The coefficient of determination measuring goodness-of-fit (r^2^) was moderately strong and ranged between 0.619-0.642 for Subject 1 and 2, respectively. Taxa that lie consistently to the left or right of the NCM-predicted relative abundance-frequency curve are those which may display some selective advantage or disadvantage, respectively. Importantly, several microbial taxa consistently deviated from the predictions of the neutral model across metacommunities, occurring less frequently than expected under neutral predictions. For instance, multiple Operational Taxonomic Units (OTUs) associated with the genera *Prevotella* and *Corynebacterium* were found to the right of the NCM distribution in the 2 subjects we surveyed, suggesting these taxa are at a selective disadvantage in replacing lost taxa relative to the assumption of neutrality among taxa (**Figure 4**).

**Figure 4:**
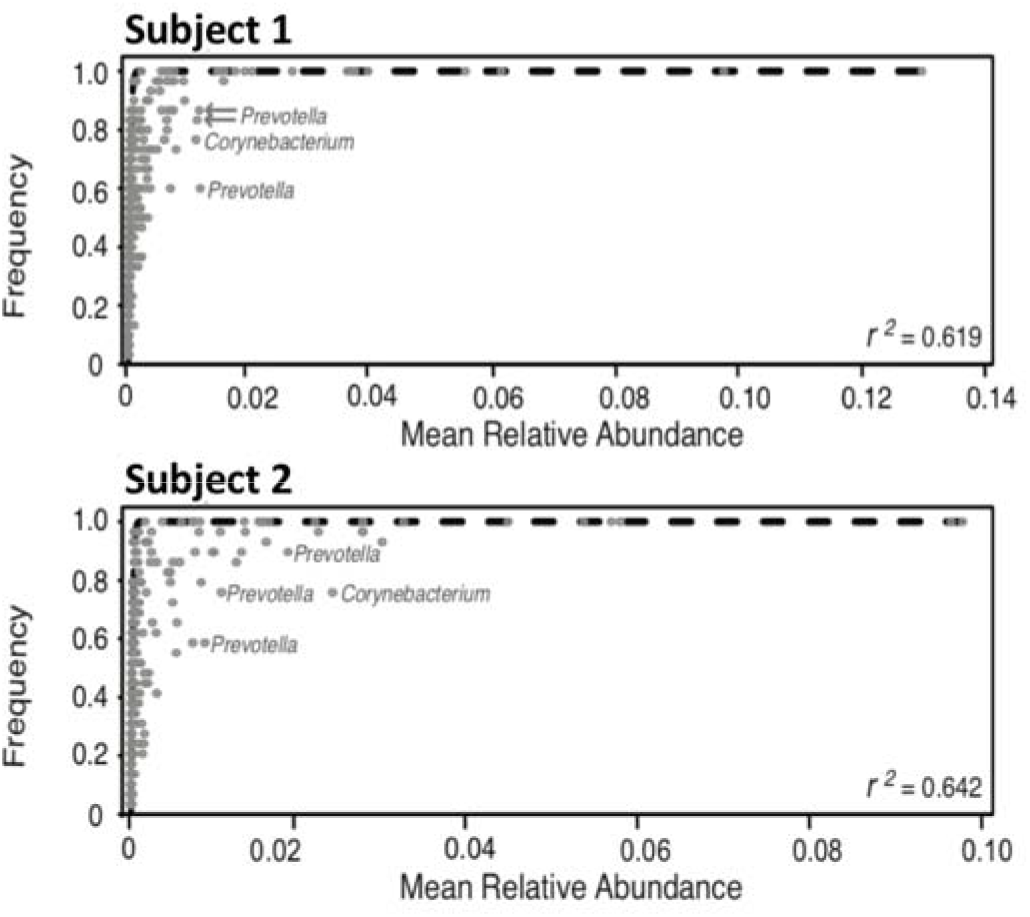
Frequency-abundance distributions of subgingival taxa conform to a neutral community model (NCM). Subject-specific comparisons of the NCM to observed frequency and mean relative abundance of subgingival OTUs. Each circle represents an individual OTU. Dashed lines represent the least-squares best fit. OTUs that lie to the right of the curve indicate taxa that may be at a selective disadvantage.

As part of the NCM we computed the migration parameter (m), which is defined as the probability that an immigrant species from a source community replaces a deceased species in the subgingival crevice. The observed overall migration parameter for each subject was small and ranged between 0.152 and 0.320 for Subject 1 and 2, respectively. These estimates are roughly on par with the migration parameters for isolated microbial communities, such as those in sewage treatment plants (m=0.1) and in the respiratory tract (m=0.2)^105^. However, they are significantly less than that previously reported for 10 control subjects and 21 patients with periodontitis for whom up to 2 samples per subject had been sequenced^110^. These disparate results may be due to differences in sequencing platform (454-Titanium in Chen et al. vs. Illumina here) or in the number of samples collected per subject. Our sample set included multiple samples of some teeth and independent samples of all teeth but the third molars, i.e. 30-31 samples per subject.

The relatively small migration parameter in this case study suggests that the majority of ‘deaths’ at each subgingival site are likely replaced through biotic reproduction of members in the local community rather than from immigration from a “mainland” source. However, a good fit to the NCM may be explained by dispersal of organisms among sites, which acts to counterbalance competitive interactions. This case study is not sufficiently powered to disentangle these possible explanations, which may serve as the focal point of further investigation.

#### Dispersal

Two models of dispersal may apply to the oral ecosystem. The first is the mainland-island model, and it assumes a near constant and relatively rapid rate of dispersal from a large source mainland to one or more islands^11^. The human oral cavity is primarily seeded from the mother and other caregivers during infancy and continues to be seeded by them soon after the eruption of the primary dentition^24^. Thus, the mainland-island model may have some bearing on the colonization of the pristine oral cavity. The parent, other caretaker, and/or older sibling may be considered as the “mainland” while the “island” or “sink” would be the infant. The infant oral cavity tends to be colonized by streptococci over the first 3 months of life^111^ with continued development of intra-oral complexity over the first year of life^112^. Factors commonly considered in mainland-island models seem to play a role in the colonization dynamics of the infant oral cavity. These factors include the bacterial load of childcare attendants (i.e., the population size of the source), the presence or absence of the primary dentition (i.e., the quality of the habitat) and the frequency of bacterial exposure^24^. For example, after primary teeth have erupted, *Streptococcus mutans*^113^ and *Aggregatibacter actinomycetemcomitans*^114^ where present in caregivers can undergo intrafamilial transmission to the infant. Understanding these dynamics may have important implications for understanding the lifetime risk of any individual to future dental disease.

The “metacommunity” island model assumes dispersal among a set of islands lacking a mainland^115^. The metacommunity model may provide insight into the transmission of disease from an affected to a non-affected tooth within an individual mouth or from saliva to a nonaffected tooth. Prior work suggests that a sound tooth surface that is adjacent to another sound tooth surface is less at risk of developing dental caries than is a tooth neighboring a carious site^116,117^. Moreover, having a filled surface is one of the strongest predictors of caries risk in individuals lacking enamel or dentinal caries while having a filled surface plus an incipient lesion is one of the strongest predictors of future caries in high individuals^118^. These observations may suggest an increase in the transmission of caries-associated bacteria from impacted to non-impacted sites. The metacommunity theory is an intriguing hypothesis worth testing while considering the alternative but complementary hypothesis that these patterns can be explained by the surface properties of different dental restorations^119^ and/or tooth morphology^120^.

A Mantel’s test can be used to test the hypothesis that geographic distance predicts similarity in the composition of bacterial communities across sites^121^. This model assumes that the relationship between the geographic distance and community dissimilarity is linear and that small or large values in one matrix correspond to similarly sized values in the other^122^. For this reason, the Mantel test can assess the extent to which dispersal, birth, death, and other contagious biotic processes induce spatial autocorrelation in the data (e.g., communities are more similar when they are geographic neighbors with incremental gains in community dissimilarity occurring with increasing geographic separation).

##### Case Study – applying the metacommunity model to the subgingival crevice

Although the NCM model suggested a low probability that an immigrant taxon from a source community replaces a deceased individual (taxon) (**Figure 4**), migration may still be high enough that inferior competitors are continuously being exchanged within the metacommunity allowing community composition to appear neutral over all. Here, we sought to test the metacommunity model using a Mantel’s test. Specifically, we compare two different models of geographic distance (island hopping vs. Euclidean distance) and Bray-Curtis dissimilarity to ask whether dispersal shapes community composition across sites in the subgingival crevice.

We refer to the first metacommunity approach that we employ as an “island-hopping” model.

The geographic distance between teeth is defined as the number of teeth separating two sample sites. Thus, the island-hopping model assumes that bacteria can only migrate step-wise from one subgingival site to an adjacent site in the same jaw. We computed the Mantel’s test for each subject independently (**Table 1**). An extremely modest, negative correlation coefficient was observed for both subjects (R^2^=-0.0376 to −0.0424) and p-values for both subjects (p>0.6) indicated that the island-hopping model is a poor fit for the subgingival crevice. In other words, a microbe that disperses from tooth 9 to tooth 15 is unlikely to traverse over tooth 10, 11, 12, 13, and 14 to reach tooth 15.

**Table 1:** Correlation between subgingival community dissimilarity and geographic distance between sites. Mantel’s test was used to examine the correlation between each geographic distance model (island-hopping, Euclidean distance) and community dissimilarity. *R* = Pearson product-moment correlation coefficient. p-values were computed after 9999 permutations and were corrected for multiple testing using the false discovery rate method.

| Subject | Dispersal model | $R^2$ | p-value |
| --- | --- | --- | --- |
| Subject 1 | **Island-hopping** | **-0.0424** | **0.651** |
|  | Euclidean distance | 0.1029 | 0.077 |
| Subject 2 | **Island-hopping** | **-0.0376** | **0.6501** |
|  | Euclidean distance | 0.0936 | 0.0869 |

The second geographic distance model assumed that the likelihood of migration from one site to another is simply a function of the direct Euclidean distance (i.e., the straight line distance) between two sites. The geographic coordinates of each sample site were obtained with 3 independent observations using a Boley gauge on a typodont model^93^. This model assumes that there are no anatomical barriers to dispersal. The correlations between Euclidean distance and Bray-Curtis dissimilarity were modest (R^2^ =0.0936-0.1029) and marginally significant (p=0.07-0.09).

Given that these data were from only 2 individuals we cannot exclude the possibility that migration and dispersal play a strong role in shaping community structure across sites in the subgingival crevice. Future work should validate these findings in a larger cohort. In addition, it is possible to use these methods to examine spatial variation at a finer scale, such as comparing communities sampled at different aspects of one tooth to those on neighboring or distal teeth. Dispersal may be hypothesized to play a role in structuring communities on a per-tooth basis, such that variation across tooth aspects on an individual tooth would appear to be less than inter-tooth variation.

### Conclusions

Consistent patterns of dental decay, gingivitis and periodontitis can be seen across human populations. That is, certain teeth and tooth aspects tend to experience higher rates of caries attack than others.^2–4,71^. Despite the spatial patterning observed in dental disease, which has been associated with local microbial communities, relatively few studies have sought to examine the spatial patterning of the human oral microbiota at mouth-wide spatial scales. Study designs that permit the collection of samples for all tooth sites and tooth aspects will enable explicit modeling of differences in community composition which may underlie population-level patterns of dental disease. We summarize what is currently known about the spatial patterning of dental disease and the associated microbiota. In addition, several statistical and ecological models were presented that can help researchers identify specific spatial patterns in the oral microbiota and the processes that underlying those patterns.

Using a case study approach, we examined the broad scale spatial patterning of the subgingival microbiota in 2 dentally healthy individuals. Through our case study, we identified an anterior-to-posterior gradient in subgingival community composition. Of particular interest, this gradient appeared to be more pronounced at mandibular compared to maxillary sites. Though the magnitude of this gradient differed by subject it was present in both individuals studied. The detection of this gradient supports our prior conclusions that some large-scale variable structures communities in the healthy human oral cavity^16^. While findings from this case study are by definition preliminary, our focus emphasizes the ways in which applying the lens of biogeography to the study of the oral microbiome will increase our understanding of it.

The development of more ecologically-informed clinical interventions for the maintenance of oral health, including the treatment of periodontitis, will not only require an understanding of biogeographic patterns, but also of the underlying ecological mechanisms that shape these patterns. From a practical perspective, while frequent sampling of all teeth and seemingly complex statistics may be important for establishing general biogeographic patterns in health and disease, it may not be necessary for detecting or managing disease in any given subject. A baseline full mouth survey and then repeated longitudinal sampling at key, risky sites may suffice for most clinical applications. The increasingly powerful, inexpensive, and pre-packaged technologies and computational algorithms required for office-based microbial community surveys in patients mean that this vision may become a reality sooner than researchers and clinicians might have imagined not so long ago.

### Data availability

A description of all methods in addition to the R code and data that were used to generate these findings can also be found at: (https://purl.stanford.edu/fx440fg9601). Raw sequencing data for this work were deposited into SRA with Qiita^123^.

### Authorship

KMS, PML, GCA, and DAR contributed to the design of IRB protocols. RK and AG analyzed samples and generated data. DMP, KMS, SPH, RK, and AG contributed to data analysis. DMP wrote the first draft of the manuscript which was revised and approved by all co-authors.

## Acknowledgements

We thank the colleagues at Stanford who provided technical assistance or instruction, including Julia Fukuyama, Elizabeth Costello and Elisabeth Bik. We thank Yvonne Kapila, the current chair at the UCSF School of Dentistry in the Department of Periodontology for departmental support. Finally, we thank the developers of Qiita and the R statistical packages used as part of this work.

## Ethical approvals

Human subjects were consented into the study in compliance with human subjects protocols approved by the University of California, San Francisco (UCSF) Human Research Protection Program and the Stanford University Administrative Panels on Human Subjects in Medical Research.

